# Novel HIV-1 fusion peptide immunogens using glycan-engineered alphavirus-like particles

**DOI:** 10.1101/2025.05.02.651772

**Authors:** Seo-Ho Oh, Dedeepya R. Gudipati, Wei Shi, Peng Zhao, Winston Wu, Jeffrey C. Boyington, Hardik K. Nariya, Emily G. McGhee, Tala Azzam, Vedhika Raghunathan, Chumeng Yang, Catherine Yang, Christian Lee, Jane D. Kim, Tongqing Zhou, John R. Mascola, Lance Wells, Rui Kong

## Abstract

Immunofocusing on conserved, subdominant epitopes is critical for vaccines against highly diverse viruses such as HIV-1, influenza, and SARS-CoV-2. The eight-residue N-terminus of the HIV-1 fusion peptide (FP) is one such example of a promising yet small target. We developed new FP immunogens using three alphavirus-like particles (VLPs) and introduced additional glycans to mask shared carrier-specific epitopes. In two independent guinea pig studies, sequential immunization with heterologous carriers enhanced FP-directed antibody titers, which were further improved with glycan engineering. Separately, using diverse FP variants sharing the same N-terminal six amino acids increased neutralizing antibody titers. When combined, these two strategies led to higher FP-directed titers and, after Env trimer boosting, induced FP-directed neutralizing antibodies against multi-clade wild-type HIV-1 in nearly all animals. These findings established the importance of minimizing recurrent off-target epitopes across immunizations and support the engineered VLPs as a promising platform for peptide immunization.

**Highlights:**

- Novel HIV-1 fusion peptide immunogens using glycan-engineered alphavirus-like particles
- Improved FP-directed response by minimizing recurrent carrier-specific epitopes across immunizations
- Improved neutralizing response by sequential immunization with diverse FP variants
- FP-directed antibodies neutralizing multi-clade wildtype viruses in nearly all animals

## Introduction

Eliciting high-titer antibody responses to conserved epitopes is critical for vaccines against highly diverse viruses^1–9^. However, these epitopes are often sub-immunodominant^10–14^. Although many advanced vaccine technologies can enhance overall immune responses^15–21^, it remains essential to maximize the on-target, subdominant response. Sequential heterologous immunization is a widely used strategy to enhance cross-reactive antibody responses against HIV-1, influenza, and SARS-Cov-2^2,6,22–34^. While most off-target epitopes present on the priming immunogen can be avoided by using boost immunogens from heterologous strains, some off-target epitopes persist and may compete with on-target epitopes for B cell engagement. For example, soluble HIV-1 Env trimers expose non-neutralizing epitopes at the trimer base and at sites where N-linked glycans are incompletely occupied^10–14,35,36^. This raises a practical question: to what extent should recurrent off-target epitopes be eliminated across sequential immunizations?

The HIV-1 fusion peptide (FP) serves as an ideal model epitope for investigating this question^37,38^, as it is a short linear sequence that can be readily distinguished from carrier-derived off-target epitopes when evaluating antibody responses. Importantly, FP is a promising target for HIV-1 vaccine development^39,40^. We previously demonstrated that immunizing non-human primates (NHPs) with an eight-amino acid FP conjugated to keyhole limpet hemocyanin (KLH), followed by booster immunizations with the HIV-1 Env trimer, elicited FP-directed antibodies capable of neutralizing up to 59% of HIV-1 strains^39^. This prime-boost strategy has since been reproducibly validated in multiple follow-up studies using varied immunization regimens and animal models^41–46^. Despite some success with conjugated vaccine approaches, it is still important to explore alternative strategies and broader rationales to enhance both the magnitude and quality of the FP-directed response prior to Env trimer boosting.

To address both the vaccinology question - to what extent recurrent off-target epitopes should be eliminated to enhance on-target antibody responses - and the specific need for high-titer FP-directed antibody responses in HIV-1 vaccine development, we developed a new set of FP immunogens using three alphavirus-like particle (VLP) carriers, including Chikungunya (CHIKV), eastern equine encephalitis (EEEV), and venezuelan equine encephalitis (VEEV) VLPs. These VLPs have been shown to be safe, well-tolerated, and highly immunogenic in clinical trials^47,48^. We also developed glycan-engineered VLPs in which off-target epitopes shared across carriers were masked. Across two independent immunization studies, FP-directed antibody responses were enhanced by using heterologous carriers in sequential immunizations and further improved when shared off-target epitopes were masked. These findings demonstrate that maximizing on-target antibody titers requires minimizing recurrent off-target epitopes across sequential immunizations. In parallel, we selected three FP variants and evaluated them in sequential immunizations, which improved the quality of the FP-directed response after three doses. When the two strategies - off-target epitope masking and heterologous FP variants - were combined, we observed a further increase in the magnitude of FP-binding and neutralizing antibodies, along with the induction of FP-directed serum antibodies capable of neutralizing multi-clade wild-type HIV-1 strains in most animals.

## Results

### HIV-1 fusion peptide immunogen design using alphavirus-like particles

CHIKV, EEEV, and VEEV VLPs are approximately 60 nm in diameter and contain 240 envelope protein E1-E2 heterodimers, which form 80 spikes^49–53^. The N-terminus of the E2 subunit is solvent-accessible, as indicated by existing VLP structures (CHIKV: PDB 6nk5^54^, EEEV: PDB 6xo4^55^, and VEEV: PDB 3j0c^53^). We designed new HIV-1 fusion peptide immunogens (CHIKV-FP8.1, EEEV-FP8.1, and VEEV-FP8.1) by genetically inserting an eight-residue FP between the E3 and E2 subunits, just C-terminal to the E3/E2 cleavage site (**Fig. 1A**). Each VLP is thus expected to present 240 FP epitopes at the N-terminus of E2 (**Fig. 1B**). Compared to unmodified VLPs, these VLP-FPs exhibited similar elution volumes in size exclusion chromatography (SEC) (**Fig. 1C**) and retained comparable size and shape in negative-stain electron microscopy (EM) (**Fig. 1D**). ELISA assays confirmed that VLP-FPs bind to both the VLP carrier-directed mAb SKT05^56^ and FP-directed mAb DFPH-a.01^39^, indicating FP epitope exposure (**Fig. 1E**). Collectively, all three VLP-FPs maintained the expected size, morphology, and antigenicity.

**Figure 1.**
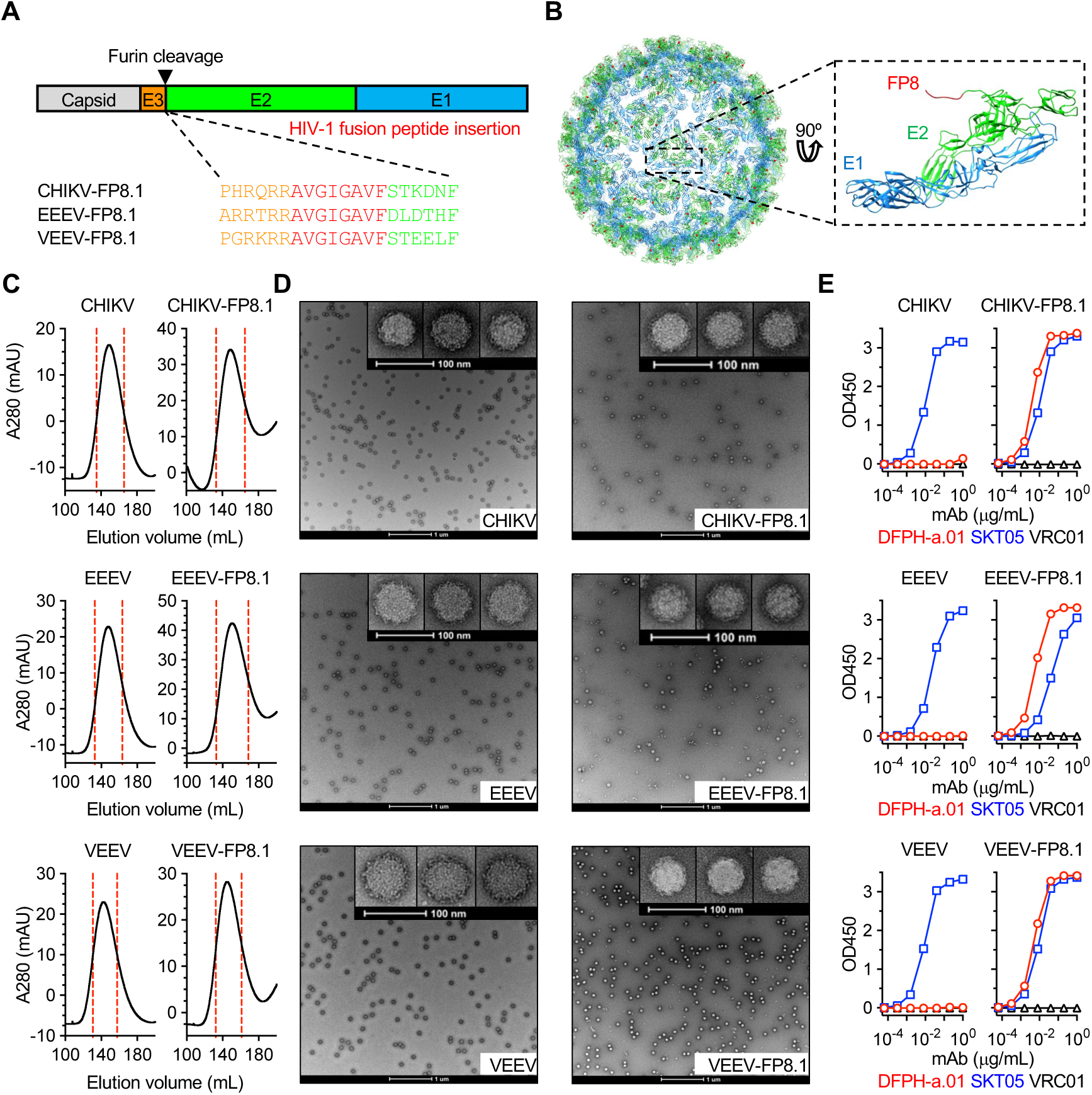
HIV-1 fusion peptide immunogen design using alphavirus-like particles. (A) Graphical schematic of the polyprotein derived from alphavirus sub-genomic RNA (termed 26S RNA) showing the HIV-1 FP insertion site between E3 and E2. (B) Ribbon diagram of the mature CHIKV T=4 icosahedral surface glycoprotein shell (left, PDB: 2XFB) and E1/E2 heterodimer (right, PDB 3N42) with FP (from PDB 6NC3) modeled at the N-terminus of E2. (C) VLP purification by size exclusion column. (D) VLP analysis by negative-stain electron microscopy. (E) VLP binding with anti-FP mAb DFPH-a.01, anti-alphavirus mAb SKT05, and anti-HIV CD4bs mAb VRC01 in ELISA.

### Glycan engineering to mask off-target epitopes shared across VLPs

To enhance the overall magnitude of FP-directed antibody response, we planned sequential FP immunizations using heterologous VLP carriers to minimize competing off-target epitopes. However, the E1 and E2 subunits of CHIKV, EEEV, and VEEV share 43-54% amino acid sequence identity (**Fig. S1**). These conserved residues (**Figs. 2A and S2**) may form off-target epitopes that compete for B cell responses during boost immunizations. To mask these shared off-target epitopes, we performed multiple iterations of glycan engineering (**Fig. 2B**), adding one glycan at a time. Using structure-based analysis^57^, we designed 50-70 potential glycan addition sites per VLP as described in the methods. Each glycan design was individually evaluated for VLP yield, FP-specific mAb binding, and VLP carrier-specific polyclonal serum binding. The top candidate was advanced to the next iteration for further glycan addition. After 3-4 iterations, we generated three new VLP-FPs: CHIKV-3g-FP8.1 (new glycans E1-69, E1-99, and E2-158), EEEV-3g-FP8.1 (new glycans E1-63, E2-28, and E2-181), and VEEV-4g-FP8.1 (new glycans E2-117, E2-150, E2-185, and E2-201) (**Figs. 2A, S3-S4**). Glycan occupancy at each of the newly introduced glycosylation sites was assessed by liquid chromatography-mass spectrometry (LC-MS) and confirmed to be high (77–99%) (**Figs. 2C and S5**).

**Figure 2.**
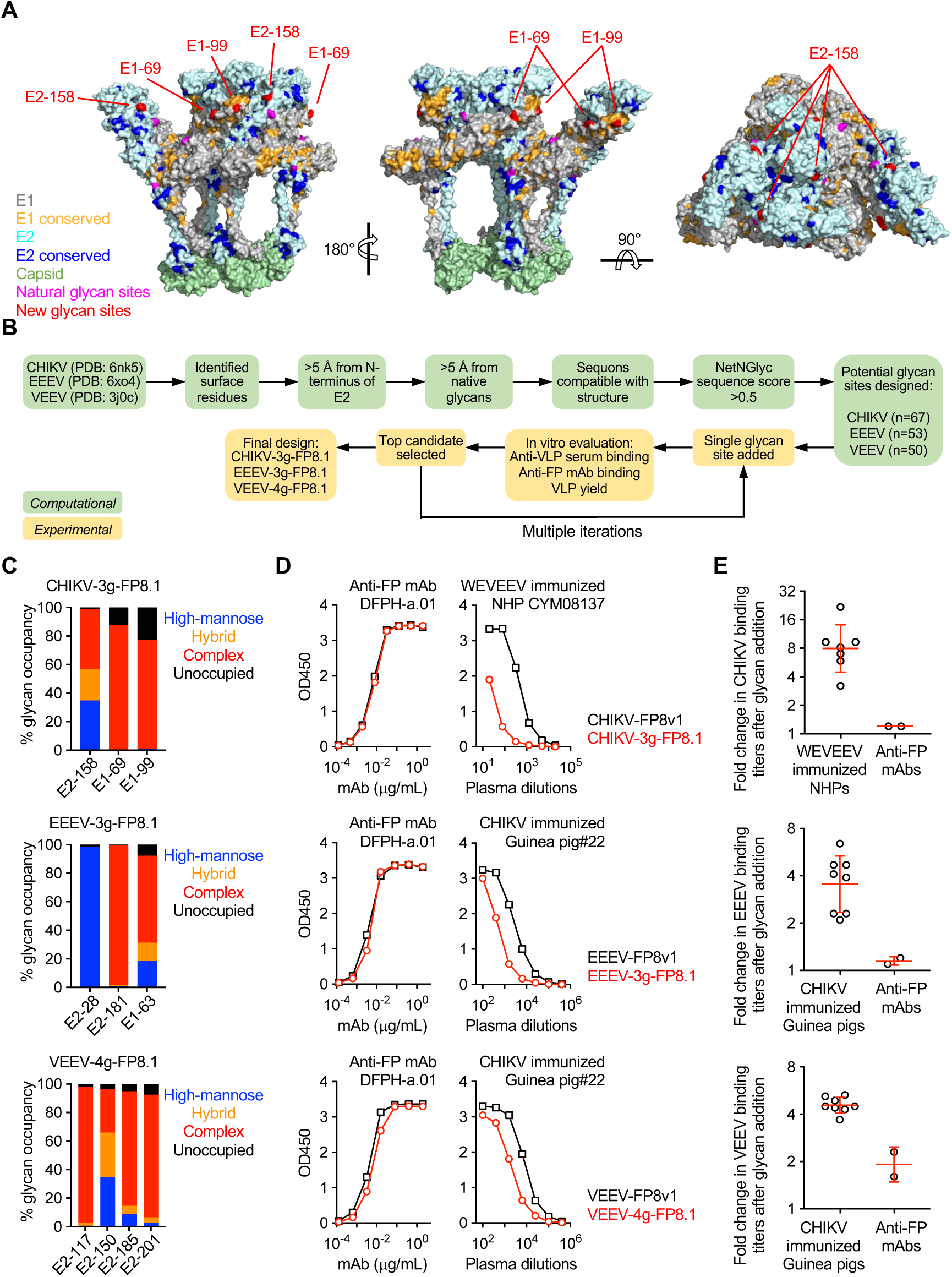
Glycan engineering to mask off-target epitopes shared across VLPs. (A) Structure of CHIKV Capsid-E1-E2 (PDB 6NK5). Conserved residues across CHIKV, EEEV, and VEEV are highlighted. Preexisting and newly added glycan sites are highlighted. (B) Flow chart illustrating the strategy for adding new glycans to the surfaces of VLPs. (C) Glycan occupancy analysis for newly added glycans. (D) mAb and animal serum binding to VLPs in ELISA. A representative animal serum is shown. WEVEEV indicates a mixture of WEEV, EEEV, and VEEV. (E) Fold change in VLP binding titers after glycan addition. VLP binding titers are defined as the serum dilution (ED1.5) or mAb concentration (EC1.5) that reach an OD450 of 1.5. Greater than one-fold change indicates reduced binding. Each circle indicates an animal serum or mAb. Red bar and error indicates geometric mean and SD. See also Figures S1-S6.

Following glycan addition, CHIKV-3g-FP8.1 VLP exhibited a geometric mean reduction of 8-fold in binding to sera from EEEV/VEEV/WEEV-immunized NHPs^56^ (**Fig. 2D and E**), indicating that about 88% of cross-reactive binding activity was blocked. Similarly, EEEV-3g-FP8.1 and VEEV-4g-FP8.1 showed geometric mean reduction of 3.5 and 4.6 fold, respectively, in binding to sera from CHIKV-immunized guinea pigs (**Fig. 2D and E**), blocking about 71% and 78% of binding activity. Importantly, glycan engineering did not affect FP-directed mAb binding (**Fig. 2D**), and the modified VLPs maintained expected profiles in SEC (**Fig. S3**) and negative-stain EM (**Fig. S4**). These results demonstrate that glycan engineering effectively masked most off-target epitopes shared across three VLP carriers without compromising particle quality or FP epitope exposure.

### Improved FP-binding antibody titers in guinea pigs after minimizing recurrent off-target epitopes across sequential immunizations

To assess the impact of sequential heterologous carrier immunization with or without glycan engineering on FP-directed antibody responses, we conducted a three-group guinea pig study (**Fig. 3A**). Group 1 received three immunizations with the same immunogen, CHIKV-FP8.1. Group 2 received three immunizations, each with a different VLP carrier. Group 3 received sequential heterologous VLP carriers with glycan engineering. Overall, Group 3 exhibited the highest FP-binding serum antibody titers after two or three immunizations (**Fig. 3B**). After two immunizations (week 6), Group 3’s geometric mean titer (70,749) was 4.3-fold higher than Group 1’s titer (16,375, P = 0.0087) and 2.2-fold higher than Group 2’s titer (32,559). Although Group 2 titers were higher than Group 1, the difference was not statistically significant. After three immunizations (week 10), Group 3’s titer (59,595) was 2-fold higher than Groups 1’s titer (29,442) and 2.2-fold higher than Group 2’s titer (27,056). These results demonstrate that while sequential heterologous carrier immunization enhances FP-directed antibody responses, minimizing recurrent off-target epitopes across immunizations is critical to maximizing this enhancement.

**Figure 3.**
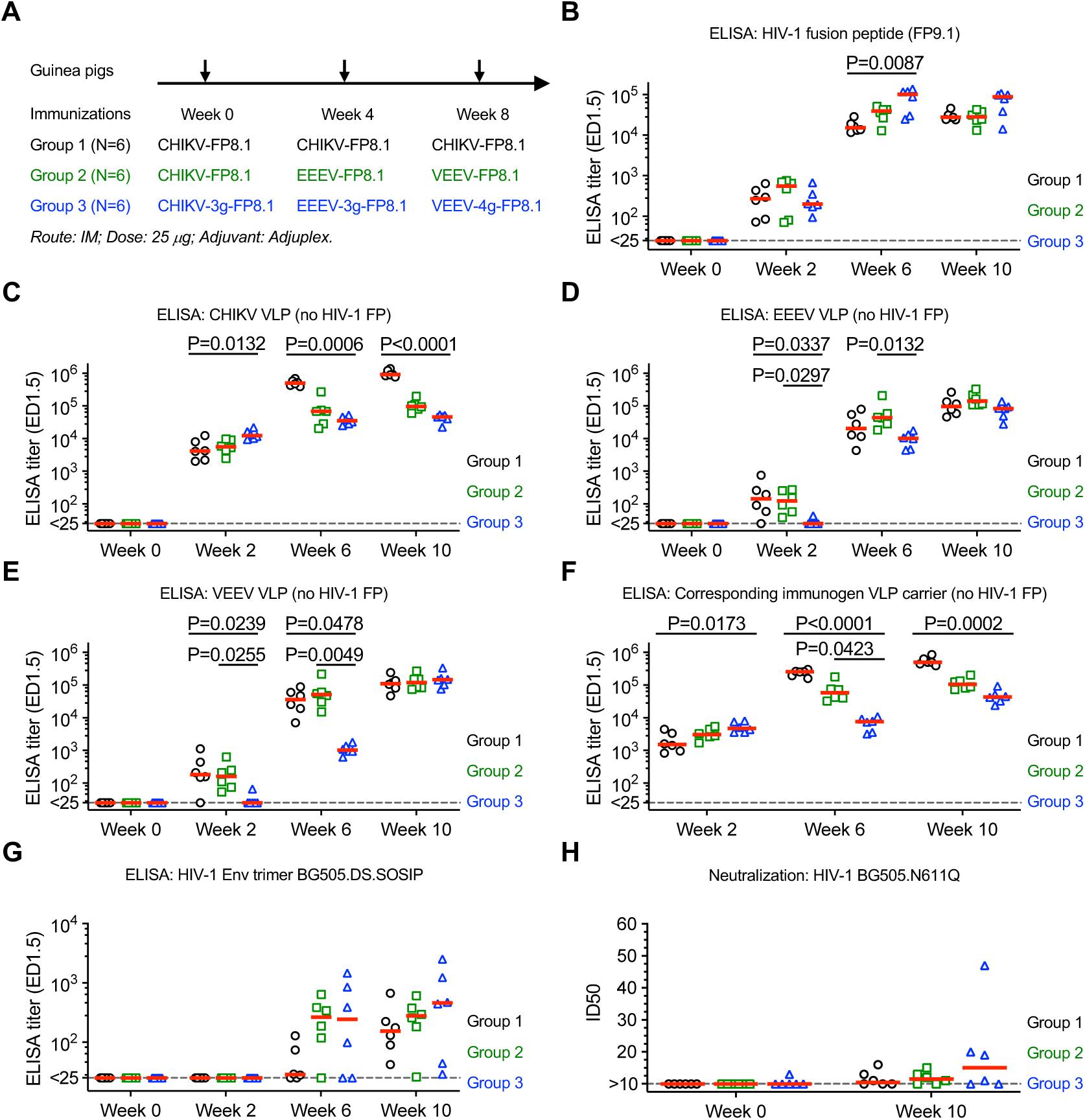
Improved FP-binding antibody titers in guinea pigs after minimizing off-target epitopes across sequential immunizations. (A) Immunization schema for guinea pig groups 1, 2 and 3. (B-G) Serum binding to free FP (panel B), CHIKV VLP (panel C), EEEV VLP (panel D), VEEV VLP (panel E), VLP carriers that match the most recent immunization (panel F), and HIV-1 Env trimer (panel G). ED1.5 indicates the serum dilution that reaches an OD450 of 1.5. (H) Serum neutralization against HIV-1 BG505.N611Q. (B-H) Median values are shown as red bar. Kruskal-Wallis test and Dunn’s multiple-comparisons test were performed. P values from significant pairs are shown.

To evaluate the impact of glycan engineering on off-target VLP-specific antibody responses, longitudinal serum samples were analyzed by ELISA for binding to unmodified CHIKV, EEEV, and VEEV VLPs (i.e., without FP insertion). Group 3 exhibited the lowest CHIKV binding after the second and third immunizations (**Fig. 3C**). Additionally, Group 3 showed little to no binding to EEEV and VEEV, whereas Groups 1 and 2 displayed detectable binding after the first immunization. This suggests that glycan engineering on CHIKV-3g-FP8.1 effectively prevented the induction of antibodies against conserved VLP carrier-specific epitopes (**Fig. 3D and E**). After the second immunization, Group 3’s geometric mean titer for EEEV binding (8,993) was 5.4-fold lower than Group 2’s titer (48,251, P = 0.0132). Similarly, Group 3’s geometric mean titer for VEEV binding (1,073) was 47.1-fold lower than Group 2’s titer (50,487, P = 0.0049). These results indicate that glycan engineering on CHIKV-3g-FP8.1 and EEEV-3g-FP8.1 effectively reduced antibody responses to conserved VLP carrier-specific epitopes (**Fig. 3D and E**). Notably, no significant difference in EEEV and VEEV binding was observed between Groups 2 and 3 after the third immunization. This is likely because glycan engineering on VEEV-4g-FP8.1 was originally designed to mask off-target epitopes shared with CHIKV (**Fig. 2**) but not specifically with EEEV. Considering that glycan engineering may introduce new epitopes not detected in ELISA using unmodified VLP carriers, we also compared antibody titers against the VLP carriers that were used in the most recent immunization (**Fig. 3F**). For example, week 6 sera from Group 3 were tested on EEEV-3g, and week 10 sera were tested on VEEV-4g. After the second immunization, Group 3’s geometric mean titer (6,363) was 36.8-fold lower than Group 1’s titer (234,474, P < 0.0001) and 9.7-fold lower than Group 2’s titer (61,783, P = 0.0423) (**Fig. 3F, week 6**). After the third immunization, Group 3’s titer (43,719) remained 12.3-fold lower than Group 1’s titer (536,759, P = 0.0002) and 2.5-fold lower than Group 2’s titer (110,243) (**Fig. 3F, week 10**). Together, these results demonstrate that glycan engineering effectively reduced antibody responses to off-target epitopes shared across CHIKV, EEEV, and VEEV VLPs. Additionally, Group 3 sera showed a trend of increased HIV-1 Env trimer-binding titers (**Fig. 3G**) and virus-neutralizing activity (**Fig. 3H**) at week 10 compared to Group 1. This suggests that enhancing the magnitude of FP-binding antibodies may also improve the magnitude of antibodies capable of recognizing FP in the context of the Env trimer.

Notably, after the first immunization (week 2), Group 3’s titers against CHIKV VLP or the corresponding immunogen carrier CHIKV-3g VLP were not lower than those of Groups 1 and 2 against CHIKV VLP (**Fig. 3C and F**), suggesting that the masked epitopes are much less dominant than other carrier-directed epitopes. Therefore, even such subdominant off-target epitopes should be eliminated across sequential immunizations to maximize the on-target antibody response.

### Improved cross-reactivity and neutralizing activity of FP-directed antibody response by sequential heterologous peptide immunizations with N-terminal six amino acid focusing

Fusion peptide sequence variation is a dominant determinant of neutralization resistance to FP-directed antibodies^37,39^. We analyzed the eight N-terminal amino acids of FP (HXB2-based numbering: 512-519) in 4736 HIV-1 sequences from the LANL database (hiv.lanl.gov) using AnalyzeAlign tool. The ten most frequent FP variants within each major HIV-1 subtype (A1, B, C, D, 01_AE or 02_AG) accounted for between ~56% and ~98% of sequences within that subtype. The top ten variants overall (FP8.1-FP8.10) covered ~57% of all 4736 sequences (**Fig. 4A**). These findings suggest that a successful FP-directed vaccine must target multiple FP variants to enhance breadth of protection. FP8.1 to FP8.10 sequence variations fall into three main categories: 1) variations at positions 518 and 519; 2) I-to-L variation at position 515; 3) T insertion at position 514b. In this study, we designed an immunization strategy (Group 4) incorporating FP8.1, FP8.2 and FP8.5 (**Fig. 4B**) and tested it alongside with Groups 1-3 described above. These three variants share the same N-terminal six amino acids but differ at positions 518 and/or 519. Rather than immunizing three times with FP8.1 (Group 2), Group 4 introduced FP8.2 in the second immunization and FP8.5 in the third. Notably, the N-terminal six amino acid focusing strategy was previously tested and recommended in prior studies^46^, where shorter peptides with seven or six amino acids were used. In contrast, our study evaluated natural eight amino acid peptide variants. After three immunizations (week 10), Group 4 exhibited a trend of improved antibody titers against FP8.1 to FP8.5 (**Fig. 4C**) compared to Group 2, though the difference was not statistically significant. This suggests that the Group 4 elicited antibodies capable of tolerating variations at positions 518 and 519 as well as L substitution at position 515. Group 4 also showed significantly higher binding titers against HIV-1 BG505.DS.SOSIP trimer (**Fig. 4D**) and significantly higher neutralizing activities against HIV-1 BG505.N611Q (**Fig. 4E**). Notably, all six animals in Group 4 developed detectable neutralizing ID50 titers, whereas most animals in Group 2 showed no neutralization. These results also suggest that the majority of FP-directed antibodies in Group 2 animals, following repeated immunizations with FP8.1, were not neutralizing. Since the BG505 trimer and virus contain the FP8.1 sequence, the improved response in Group 4 is likely due to N-terminal six amino acid focusing.

**Figure 4.**
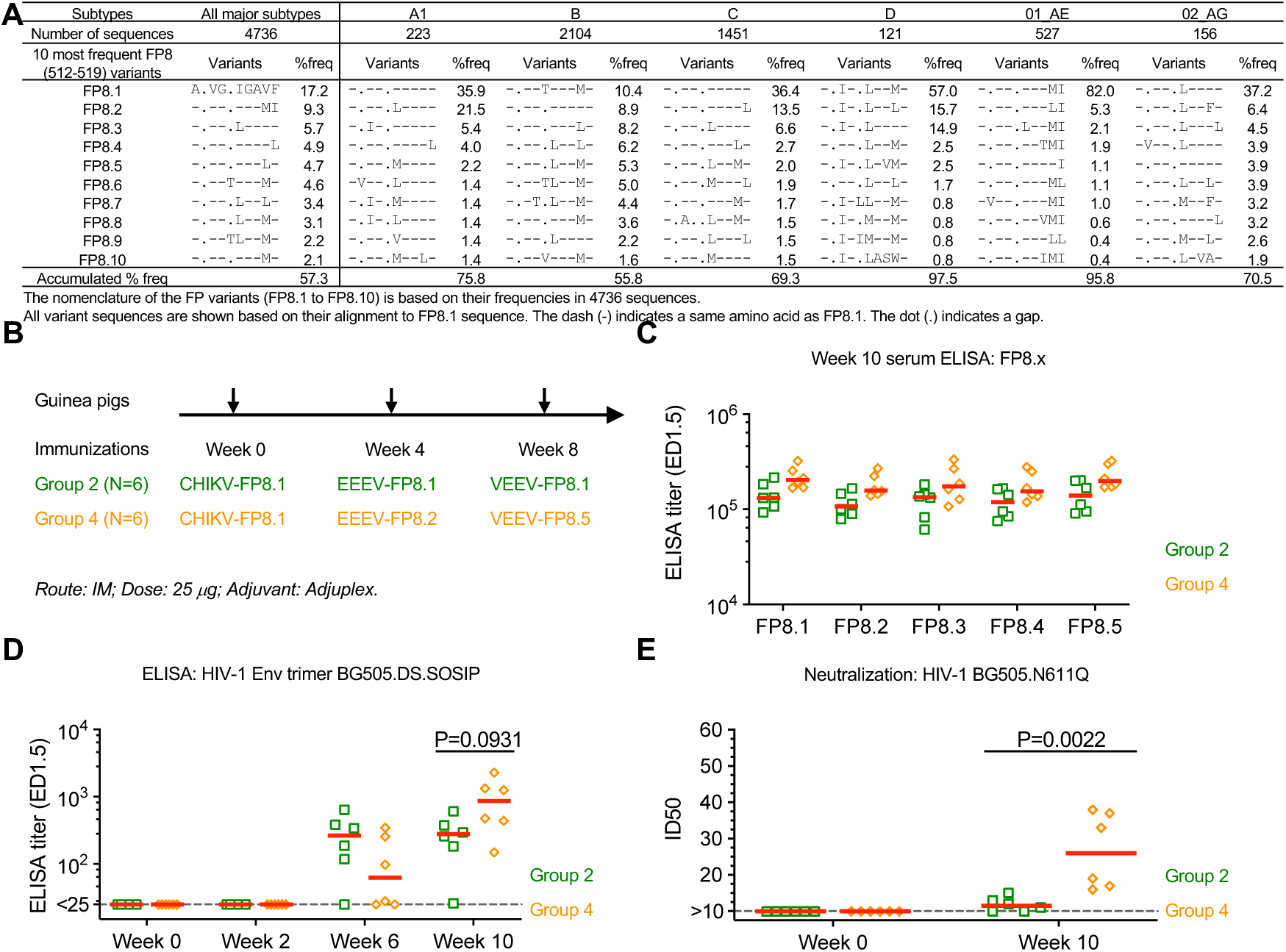
Improved cross-reactivity and neutralizing activity of FP-directed antibody response by sequential heterologous peptide immunizations with N-terminal six amino acid focusing. (A) Analysis of FP variants and their frequencies in 4736 HIV-1 sequences from the LANL database. (B) Immunization schema for guinea pig groups 2 and 4. (C-D) Serum binding to free FP8 variants (panel C) and HIV-1 Env trimer (panel D). ED1.5 indicates the serum dilution that reaches an OD450 of 1.5. (E) Serum neutralization against HIV-1 BG505.N611Q. (C-E) Median values are shown as red bar. Kruskal-Wallis test and Dunn’s multiple-comparisons test were performed. P values from significant pairs are shown.

### Further enhancement of FP-directed binding and neutralizing antibody response through a combination of two strategies

Since Group 3 and 4 independently improved FP-directed antibody response through distinct mechanisms, we conducted a Group 5 study (**Fig. 5A**) to evaluate the combined effects of sequential heterologous VLP carrier with glycan engineering and heterologous peptides with N-terminal focusing. Group 5 animals received sequential immunizations with CHIKV-3g-FP8.1, EEEV-3g-FP8.2, and VEEV-4g-FP8.5. Meanwhile, two control groups were included: Group 6 received three immunizations with the same CHIKV-3g carrier. Group 7, replicating Group 4, received sequential immunizations with CHIKV-FP8.1, EEEV-FP8.2, and VEEV-FP8.5.

**Figure 5.**
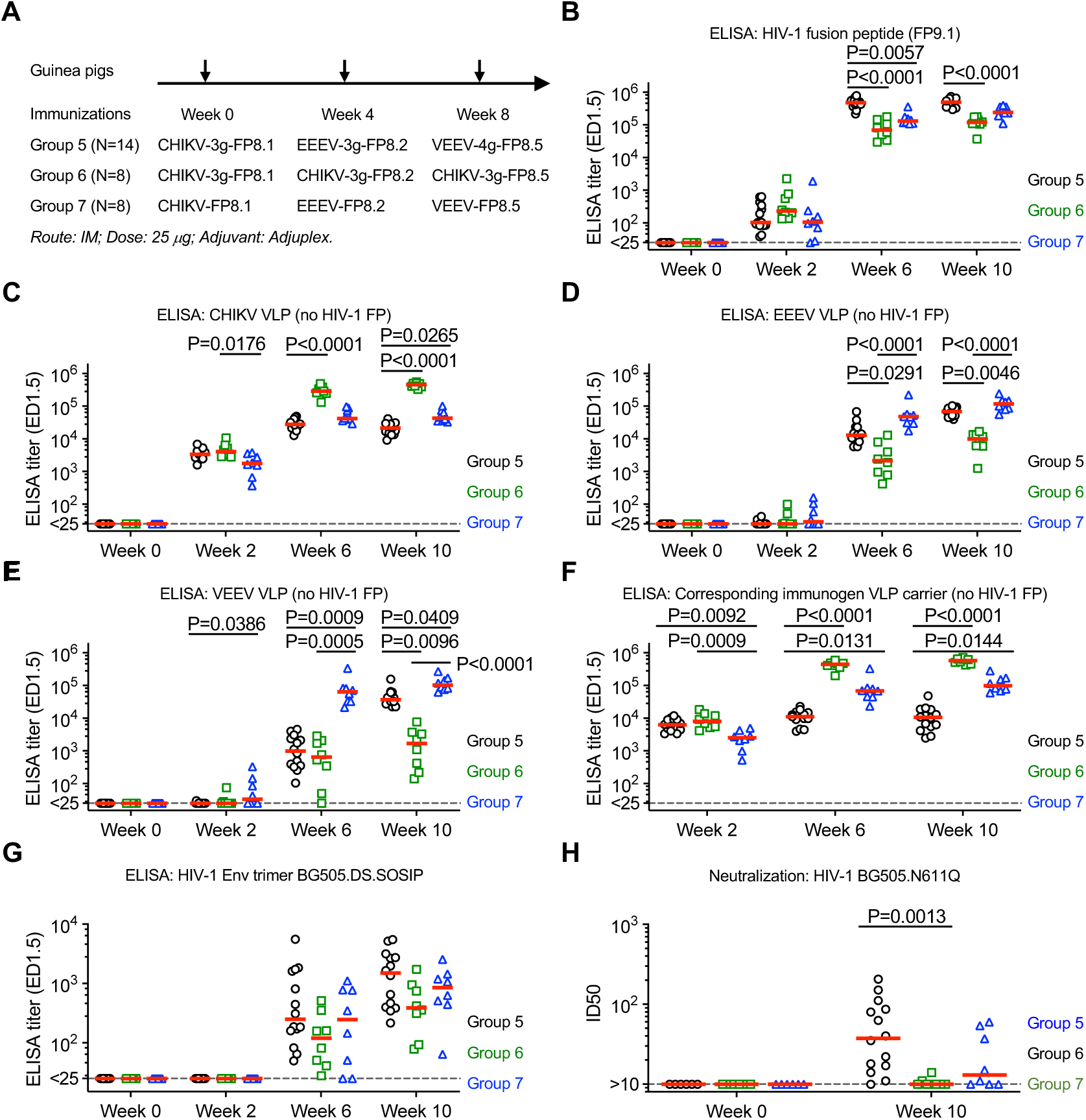
Further enhancement of FP-directed binding and neutralizing antibody response through a combination of two strategies. (A) VLP immunization schema for guinea pig groups 5-7. (B-G) Serum binding to free FP (panel B), CHIKV VLP (panel C), EEEV VLP (panel D), VEEV VLP (panel E), VLP carriers that match the most recent immunization (panel F), and HIV-1 Env trimer (panel G). ED1.5 indicates the serum dilution that reaches an OD450 of 1.5. (H) Serum neutralization against HIV-1 BG505.N611Q. (B-H) Median values are shown as red bar. Kruskal-Wallis test and Dunn’s multiple-comparisons test were performed. P values from significant pairs are shown.

Overall, Group 5 exhibited the highest FP-binding serum antibody titers after second and third immunizations (**Fig. 5B**). After two immunizations (week 6), Group 5’s geometric mean titer (461,000) was 6.4-fold higher than Group 6 (71,512, P < 0.0001) and 3.1-fold higher than Group 7 (149,355, P = 0.0057). After three immunizations (week 10), Group 5’s titer (462,743) was 4.1-fold higher than Group 6 (113,297, P < 0.0001) and 1.9-fold higher than Group 7 (250,007). These results again demonstrated that minimizing off-target epitopes across immunizations is critical to maximizing the on-target antibody response.

Similar to the Groups 1-3 study, Group 5 exhibited the lowest binding to the CHIKV VLP carrier (without FP insertion) after the second and third immunizations (**Fig. 5C**). After three immunizations, Group 5’s geometric mean titer (20,678) was 2.4-fold lower than Group 7’ (49,996, P = 0.0265). Additionally, Group 5’s titers for the VEEV VLP carrier were significantly lower than those in Group 7 after each immunization (Post-1: P = 0.0386; Post-2: 67.9-fold, P = 0.0009; Post-3: 2.8-fold, P = 0.0409) (**Fig. 5E**). Group 5’s titers for the EEEV VLP carrier were also lower than those in Group 7 after the second and third immunizations (Post-2: 3.3-fold; post-3: 1.7-fold), those the difference were not statistically significant (**Fig. 5D**). We further compared antibody titers against the VLP carriers used in the most recent immunization (**Fig. 5F**). After the secon immunization, Group 5’s geometric mean titer (10,261) was 40-fold lower than Group 6 (410,767, P < 0.0001) and 6.7-fold lower than Group 7 (69,020, P = 0.0131) (**Fig. 5F, week 6**). After the third immunization, Group 5’s titer (9,409) remained 59.5-fold lower than Group 6 (559,985, P < 0.0001) and 11.8-fold lower than Group 7 (111,016, P = 0.0144) (**Fig. 5F, week 10**). Together, these results are consistent with those from the Group 1-3 study, further demonstrating that glycan engineering effectively reduces antibody responses to off-target epitopes shared across CHIKV, EEEV, and VEEV VLPs. Additionally, Group 5 sera showed a trend of increased HIV-1 Env trimer-binding (**Fig. 5G**).

After three immunizations, 13 of 14 animals (93%) in Group 5 developed detectable neutralizing activity against BG505.N611Q, compared to 2 of 8 (25%) in Group 6 and 5 of 8 (63%) in Group 7. Group 5’s median ID50 titer (39.5) was 3.7-fold higher than Group 6 (10.6, P = 0.0013) and 2-fold higher than Group 7 (19.4). These results indicate that the neutralizing antibody titers were further enhanced by combining the two immunization strategies.

### FP-directed serum antibodies neutralize heterologous wildtype HIV-1 strains across multiple clades

Animals in Group 5-7 were further boosted four times with soluble HIV-1 Env trimers BG505.DS.SOSIP and ConC.DS.SOSIP (**Fig. 6A**). To evaluate serum antibody neutralizing activity, total IgG was purified from week 27 sera and tested against wildtype HIV-1 strains from multiple clades. Neutralizing antibodies were detected in most Group 5 animals (**Figs. 6B and S6**). BI369A (clade A) was neutralized by purified IgG from all animals, with a geometric mean IC50 of 1.4 mg/mL. BL01.DG (clade B) was neutralized by IgG from 86% of animals, with a geometric mean IC50 of 2.3 mg/mL. 286.36 (clade C) was neutralized by IgG from 29% of animals, with a geometric mean IC50 of 2.7 mg/mL. CH117.4 (clade BC) was neutralized by IgG from all animals, with a geometric mean IC50 of 1 mg/mL. Importantly, no non-specific neutralization was observed against the control virus SIVmac251, confirming the specificity of the response.

**Figure 6.**
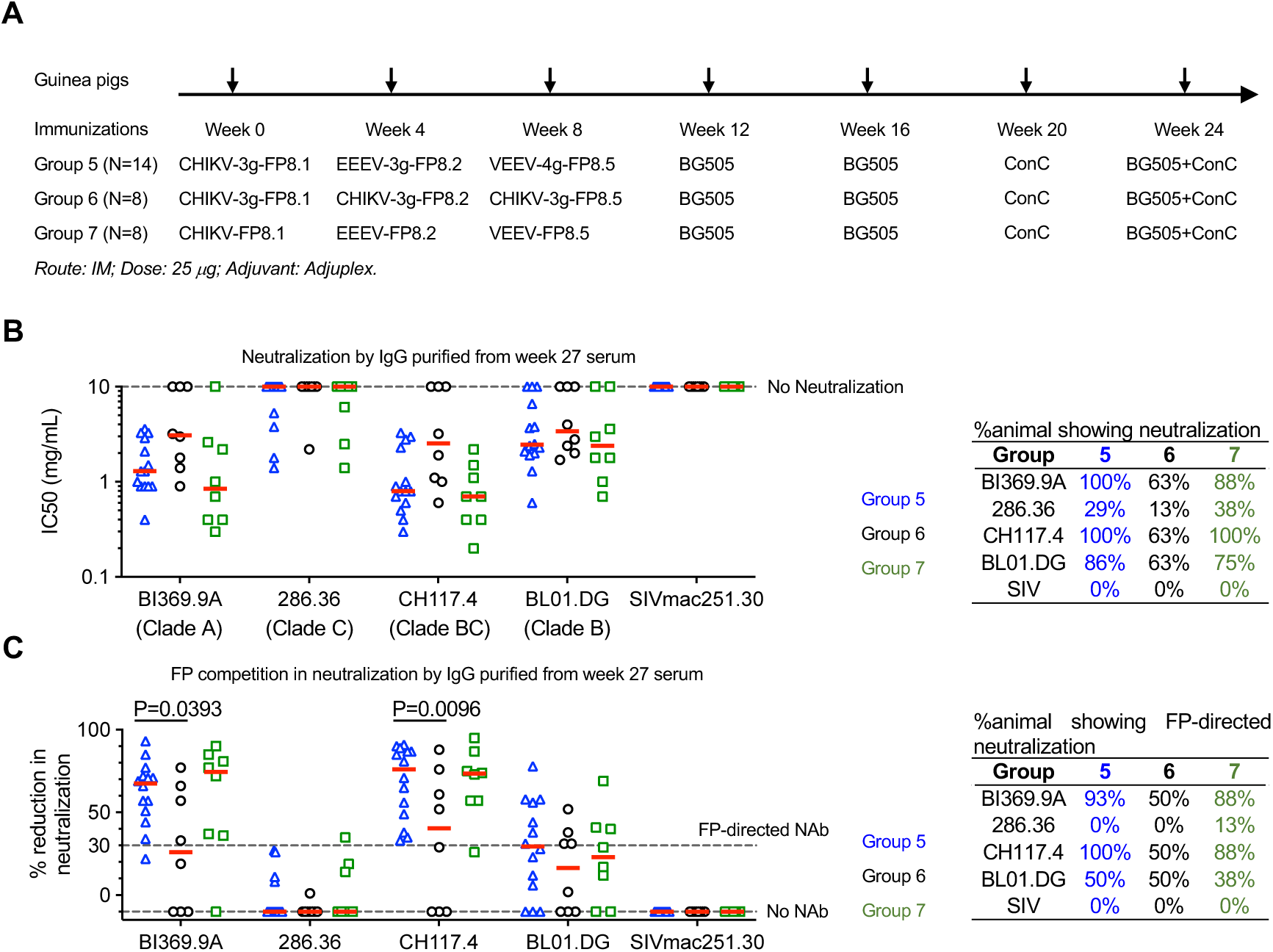
FP-directed serum antibodies neutralize heterologous wildtype HIV-1 strains from multiple clades. (A) Prime and boost immunization schema for guinea pig groups 5-7. (B) Neutralization IC50 titers of IgG purified from week 27 sera against wildtype HIV-1 strains from multiple clades and the control SIVmac251 strain. (C) % reduction in serum IgG neutralization in the presence of FP8v1 peptide. Median values are shown as red bar. Fisher’s exact test were performed. P values from significant pairs are shown.

To determine whether the observed neutralization was mediated by FP-directed antibodies, the same neutralization assays were performed in the presence of free FP peptide. IC50 titers with and without FP competition were compared, and percent reduction in neutralization was calculated. A reduction greater than 30% was defined as evidence of FP-directed neutralizing activity. Most of the neutralization observed in Group 5 was FP-dependent (**Figs. 6C and S6**). About 93% of samples showed FP-directed neutralization against BI369.9A, with an average 65% reduction in neutralization. About 50% of sample showed FP-directed neutralization against BL01.DG, with an average 52% reduction. All sample showed FP-directed neutralization against CH117.4, with an average 68% reduction. No FP-directed neutralization was observed against 286.36. Together, these results demonstrate that FP-directed antibodies capable of neutralizing multi-clade wildtype strains were elicited as a dominant component of the antibody response in most Group 5 animals.

Notably, FP-directed neutralizing antibodies were also detected in a substantial proportion of Group 6 and 7 animals (**Figs. 6B-C and S6**). However, the percent reduction in neutralization against BI369.9A and CH117.4 was significantly lower in Group 6 compared to Group 5 (**Fig. 6C**). This finding highlights the advantage of sequential heterologous carrier immunization over repetitive immunization. At the same time, it suggests that further optimization of trimer boost strategies may be necessary to fully reveal or amplify the benefits conferred by the optimized VLP immunization regimen.

## Discussion

Sequential heterologous immunization is a widely used strategy for targeting conserved, yet sub-immunodominant, epitopes in highly diverse viruses such as HIV-1^2^, influenza^6,22–28^, and SARS-CoV-2^29–31^. However, when designing and optimizing boost immunogens, it remains unclear to what extent shared off-target epitopes should be eliminated to maximize the on-target antibody response^10–14,35,36^. Here we investigated the impact of masking recurrent off-target epitopes across immunizations on the on-target antibody response in the context of sequential HIV-1 FP immunizations. We generated new FP immunogens by genetically inserting the FP sequence at the N-terminus of the E2 subunits of CHIKV, EEEV, and VEEV VLPs (**Fig. 1**) and engineered additional glycans to mask carrier-directed epitopes shared across the three VLPs (**Fig. 2**). When CHIKV, EEEV, and VEEV VLP carriers were used sequentially, we anticipated that off-target B cells recognizing shared carrier epitopes would be boosted and could compete with FP-specific B cells. In two independent guinea pig studies, CHIKV-FP8.1 VLP induced antibodies that cross-react with EEEV and VEEV VLPs (**Fig. 3D and E**, week 2; **Fig. 5D and E**, week 2). In the same two studies, following the second immunization, EEEV-3g-FP8.1—with added glycans—showed reduced serum binding to the EEEV carrier by 5.4-and 3.3-fold compared to EEEV-FP8.1 (**Figs. 3D and 5D**, week 6), suggesting that shared carrier-directed epitopes contributed substantially to the total off-target response. Concurrently, FP-directed antibody titers increased (**Figs. 3B and 5B**, week 6). These results demonstrate that reducing recurrent off-target epitopes across sequential heterologous immunizations can further enhance the absolute titers of the on-target FP-directed antibody response.

Since inducing high-titer antibody responses to highly conserved viral epitopes is critical for broadly protective vaccine development^1–9^, the results above recommend to minimize off-target epitopes shared across sequential heterologous immunizations. For example, sequential immunization with HIV-1 soluble trimers is currently under extensive evaluation in animal models^2,32–34,58^ and clinical trials^59–62^. However, previous studies have shown that these trimers often elicit dominant off-target responses to non-neutralizing epitopes^10–14^, particularly at the artificially exposed trimer base. While efforts to address these base epitopes have begun^14,35^, immunogen designs that eliminate base-directed responses and enhance on-target immunogenicity have not yet been established in vivo^14,35^, and it remained unclear whether such intervention was necessary. Our findings suggest that appropriately addressing off-target base epitopes may significantly enhance antibody responses to broadly reactive neutralizing epitopes, providing a strong rationale for their removal in future immunogen designs.

The sequential immunization strategy using heterologous alphavirus VLP carriers with glycan engineering, when combined with heterologous natural FP variants incorporating N-terminal six–amino acid focusing, further enhanced the magnitude of FP-directed binding (**Fig. 5B**) and neutralizing antibody titers (**Fig. 5H**). Importantly, following trimer boost immunizations, nearly all animals developed FP-directed antibodies capable of neutralizing heterologous wildtype viruses from multiple HIV-1 clades (**Fig. 6**). Given that the unmodified parental VLPs have been shown to be safe and well tolerated in previous clinical trials^47,48,50^, these VLP-FP designs represent promising candidate components for future FP-directed HIV-1 vaccine regimens. Notably, serum neutralization titers against wildtype viruses still require improvement. Potential strategies could include: (1) next-generation VLP designs with minimized total off-target epitopes, not limited to those shared across VLPs; (2) application of novel multivalent immunogen platforms with minimal off-target immunogenicity, such as DNA origami scaffolds^63,64^; (3) FP epitope optimization to better mimic its native presentation in the Env trimer; and (4) improved Env trimer designs for boost immunizations.

Finally, these glycan-engineered VLP designs may have broader utility and could be adapted to present key epitopes from other pathogens or therapeutic targets.

## Acknowledgments

We thank M. Roederer for contributing serum samples from NHPs immunized with EEEV, VEEV, and WEEV VLPs. We thank Vaccine Production Program of NIAID VRC for contributing adjuvant Adjuplex. We thank Y. Li for sharing serum IgG purification experience. Support for this work was provided by the NIH (R01AI162267 to R.K. and the intramural research program to J.R.M), the NSF Biofoundry (Glycoscience Research, Education, and Training Grant 2400220 to L.W.), Emory Vaccine Center (R.K.), and Georgia Research Alliance (R.K.). Emory National Primate Research Center (ENPRC) was supported by NIH Office of Research Infrastructure Programs (ORIP) P51 OD11132.

## Author Contributions

S.O., H.K.N. and E.G.M. performed immunization studies and analysis; D.R.G., T.A., V.R., Chumeng Y., Catherine Y. and R.K. developed glycan engineering of VLPs; W.S., W.W, J.C.B., J.R.M, and R.K. designed and tested the original CHIKV-FP8.1 VLP; P.Z. and L.W. performed glycan analysis; S.O., H.K.N., and C.L. produced VLPs and performed EM analysis; T.Z. provided trimer proteins.

## STAR★METHODS

### RESOURCE AVAILABILITY

#### Lead contact

Further information and requests for resources and reagents should be directed to and will be fulfilled by the lead contact, Rui Kong.

#### Materials availability

Materials used in this study are available upon request to the lead contact.

#### Data and code availability

The mass spectrometry data have been deposited to the MassIVE database (https://massive.ucsd.edu/ProteoSAFe/static/massive.jsp) with the identifier MSV000097434 (ftp://MSV000097434@massive.ucsd.edu). Any additional information required to reanalyze data reported in this paper is available from the lead contact upon request.

### EXPERIMENTAL MODEL AND SUBJECT DETAILS

#### Guinea pig

Female Hartley guinea pigs with body weights of 300-350 g each were purchased from Charles River Laboratories. Guinea pigs were housed and cared for in accordance with local, state, federal, and institute policies in an Association for Assessment and Accreditation of Laboratory Animal Care International (AAALAC)-accredited facility at the Emory National Primate Research Center. The animal experiments were reviewed and approved by the Institutional Animal Care and Use Committee of Emory University and covered under protocol PROTO202000164.

#### Cell lines

Expi293F cells were from ThermoFisher Scientific Inc (Invitrogen, Cat# A14528; RRID: CVCL_D615). HEK293T/17 cells were from ATCC (Cat# CRL-11268). TZM-bl cells were from NIH AIDS Reagent Program (www.aidsreagent.org, Cat# 8129). The cell lines were cultured following manufacturer suggestions as described in Method Details below.

### METHOD DETAILS

#### Virus-like particle production

Expi293F cells were maintained in Expi293 Expression Medium (Gibco, A14351-01) at 37°C with 8% CO₂ in a shaking incubator at 125 rpm. Cells were seeded at a density of 2.5 × 10⁶ cells/mL and transfected with a plasmid encoding the structural proteins of the alphavirus using linear 25 kDa polyethylenimine (PEI) (Polysciences, Cat# 23966-1). The following day, enhancers were added to the culture to boost protein expression. After five additional days, culture supernatant was harvested, filtered and concentrated by tangential flow filtration (TFF). The concentrated supernatant was mixed 1:1 (v/v) with OptiPrep™ (60% iodixanol solution; ThermoFisher Scientific, NC1059560), transferred into Quick-Seal centrifuge tubes (Beckman Coulter, Cat# 344322), sealed, and ultracentrifuged at 70,000 rpm for 11 hours at 4 °C using a Beckman Coulter Type 70.1 Ti rotor. After centrifugation, VLP-containing fraction was collected, filtered through a 0.45 µm membrane, and further purified by size exclusion chromatography (SEC) using a HiPrep™ 26/60 Sephacryl™ S-500 HR column (Cytiva, 28-9356-07) on an ÄKTA Pure system (Cytiva). Peak fractions were pooled and concentrated using Amicon Ultra-70 centrifugal filter units (30 kDa MWCO; Millipore Sigma, UFC703008). Purified VLPs were then filtered through a 0.2 µm membrane, aliquoted, and stored at –80 °C.

#### Negative-Stain Electron Microscopy

A 3 µL aliquot of purified VLPs (0.05–0.10 mg/mL) was applied to glow-discharged, carbon-coated 400-mesh copper grids and incubated for 60 seconds at room temperature. Excess liquid was removed by blotting with filter paper. Grids were washed twice with 10 µL of washing buffer (10 mM HEPES, 150 mM NaCl, pH 7.0), with blotting after each wash to remove residual buffer. The particles were negatively stained with 10 µL of 0.75% (w/v) uranyl formate for 60 seconds. Excess stain was blotted away, and grids were air-dried for 5 minutes at room temperature. Imaging was performed using a Thermo Fisher Talos L120C transmission electron microscope (TEM) operating at 120 kV. Images were acquired at magnifications of 11,000×, 73,000×, and 92,000× to visualize the VLPs.

#### Glycan engineering

Glycan engineering was performed as previously described^57^. Briefly, using crystal structures of CHIKV, EEEV, and VEEV VLPs, surface-exposed residues on the E1 and E2 proteins located more than 5 Å from the E2 N-terminus or existing glycosylation sites were identified. Each candidate site was examined in PyMOL (The PyMol Molecular Graphics System, version 1.8.6; Schrodinger, LLC), and positions where incorporation of an NxT sequon was unlikely to disrupt protein stability were selected. These positions were further evaluated for glycosylation potential using the NetNGlyc server (http://www.cbs.dtu.dk/services/NetNGlyc/), and those with prediction scores above 0.5 were selected for VLP mutagenesis. Individual glycan-modified VLPs were expressed and assessed for yield, anti-FP monoclonal antibody binding, and serum binding. The top-performing construct was then used to evaluate the addition of further glycans, as described in the main text.

#### Analysis of site-specific N-linked glycopeptides for VLPs proteins by LC-MS

As described previously^65^, aliquots of the purified proteins were reduced by incubating with 5 mM of dithiothreitol (Sigma) at 56 °C and alkylated by 13.75 mM of iodoacetamide (Sigma) at room temperature in dark. The aliquots were then digested respectively by using a combination of alpha lytic protease (New England BioLabs), AspN (Promega), chymotrypsin (Athens Research and Technology), Lys-C (Promega), Glu-C (Promega), and trypsin (Promega). The resulting peptides were separated on an Acclaim™ PepMap™ 100 C18 column (75 µm x 15 cm) and eluted into the nano-electrospray ion source of an Orbitrap Eclipse™ Tribrid™ mass spectrometer (Thermo Scientific) at a flow rate of 200 nL/min. The elution gradient consists of 1-40% acetonitrile in 0.1% formic acid over 370 minutes followed by 10 minutes of 80% acetonitrile in 0.1% formic acid. The spray voltage was set to 2.2 kV and the temperature of the heated capillary was set to 275 °C. Full MS scans were acquired from m/z 200 to 2000 at 60k resolution, and MS/MS scans following higher-energy collisional dissociation (HCD) with stepped collision energy (15%, 25%, 35%) were collected in the orbitrap at 15k resolution. pGlyco3^66^ was used for database searches with mass tolerance set as 20 ppm for both precursors and fragments. The database search output was filtered to reach a 1% false discovery rate for glycans and 10% for peptides. The glycan and peptide assignment for each spectra was then manually validated after filtering. Quantitation was performed by calculating spectral counts for each glycan composition at each site. Any N-linked glycan compositions identified by only one spectra were removed from quantitation. N-linked glycan compositions were categorized into 19 classes (including “Unoccupied” as class 19):

HexNAc(2)Hex(9~5)Fuc(0~1) was classified as M9 to M5 respectively;

HexNAc(2)Hex(4~1)Fuc(0~1) was classified as M1-M4;

HexNAc(3~6)Hex(5~9)Fuc(0)NeuAc(0~1) was classified as Hybrid with

HexNAc(3~6)Hex(5~9)Fuc(1~2)NeuAc(0~1) classified as F-Hybrid; Complex-type glycans are classified based on the number of antenna and fucosylation:

HexNAc(3)Hex(3~4)Fuc(0)NeuAc(0~1) is assigned as A1 with

HexNAc(3)Hex(3~4)Fuc(1~2)NeuAc(0~1) assigned as F-A1;

HexNAc(4)Hex(3~5)Fuc(0)NeuAc(0~2) is assigned as A2/A1B with

HexNAc(4)Hex(3~5)Fuc(1~5)NeuAc(0~2) assigned as F-A2/A1B;

HexNAc(5)Hex(3~6)Fuc(0)NeuAc(0~3) is assigned as A3/A2B with

HexNAc(5)Hex(3~6)Fuc(1~3)NeuAc(0~3) assigned as F-A3/A2B;

HexNAc(6)Hex(3~7)Fuc(0)NeuAc(0~4) is assigned as A4/A3B with

HexNAc(6)Hex(3~7)Fuc(1~3)NeuAc(0~4) assigned as F-A4/A3B;

HexNAc(7)Hex(3~8)Fuc(0)NeuAc(0~1) is assigned as A5/A4B with

HexNAc(7)Hex(3~8)Fuc(1~3)NeuAc(0~1) assigned as F-A5/A4B.

#### Analysis of deglycosylated VLPs proteins by LC-MS

As described previously^65^, aliquots of the purified proteins were reduced by incubating with 5 mM of dithiothreitol (Sigma) at 56 °C and alkylated by 13.75 mM of iodoacetamide (Sigma) at room temperature in dark. The aliquots were then digested respectively using chymotrypsin (Athens Research and Technology), Glu-C (Promega), Asp-N (Promega), and trypsin (Promega). Following digestion, the extracted peptides were deglycosylated by Endoglycosidase H (Promega) followed by PNGaseF (Promega) treatment in the presence of 18O water (Cambridge Isotope Laboratories). The resulting peptides were separated on an Acclaim™ PepMap™ 100 C18 column (75 µm x 15 cm) and eluted into the nano-electrospray ion source of an Orbitrap Eclipse™ Tribrid™ mass spectrometer (Thermo Scientific) at a flow rate of 200 nL/min. The elution gradient consists of 1-40% acetonitrile in 0.1% formic acid over 370 minutes followed by 10 minutes of 80% acetonitrile in 0.1% formic acid. The spray voltage was set to 2.2 kV and the temperature of the heated capillary was set to 275 °C. Full MS scans were acquired from m/z 200 to 2000 at 60k resolution, and MS/MS scans following higher-energy collisional dissociation (HCD) with stepped collision energy (15%, 25%, 35%) were collected in the ion trap. The spectra were analyzed using SEQUEST (Proteome Discoverer 2.5, Thermo Fisher Scientific) as well as Byonic (v4.1.10, Protein Metrics) with mass tolerance set as 20 ppm for precursors and 0.5 Da for fragments. The search output was filtered to reach a 1% false discovery rate at the protein level and 10% at the peptide level. The site assignment for each spectra was then manually validated after filtering. Occupancy of each N-linked glycosylation site was calculated using spectral counts assigned to the 18O-Asp-containing (PNGaseF-cleaved) and/or HexNAc-modified (EndoH-cleaved) peptides and their unmodified counterparts.

#### Quantification and statistical analysis

Raw glycoproteomic data from the mass spectrometers was searched using SEQUEST (Proteome Discoverer 2.5), Byonic (v4.1.10), and pGlyco3^66^. Search results from SEQUEST and Byonic were filtered to reach a 1% false discovery rate at the protein level and 10% at the peptide level; and search results from pGlyco3 was filtered to reach a 1% false discovery rate at the glycan level and 10% at the peptide level. All spectral assignments were manually validated after applying false discovery rate filtering.

#### Animal experiment

Female Hartley guinea pigs (300–350 g) were obtained from Charles River Laboratories. Following a two-week acclimation period, animals were grouped to ensure comparable mean body weights prior to immunization. Animals were immunized at 4-week intervals, and sera were collected at week 0 and two weeks after each immunization. For each immunization, 400 µL of immunogen mixture containing 25 µg of VLP or trimer immunogen and 80 µL of Adjuplex (obtained from VRC, NIAID, NIH) was administered evenly into the quadriceps muscles of both hind legs. For serum collection, blood was drawn via saphenous vein bleeding into serum separator tubes and centrifuged to isolate serum. Serum samples were aliquoted, snap-frozen in liquid nitrogen, and stored at –80 °C until further analysis.

#### Serum IgG purification

The Serum IgG was purified by AmMag™ Protein A Magnetic Beads (Genscript Biotech, L00695). 1 mL of serum was diluted into 9 mL of PBS and mixed with 250 μL of AmMag^TM^ Protein A Magnetic Beads in 1 mL of 20% ethanol slurry in a 15 mL conical tube. Then, the tube was incubated at room temperature with mixing on a rocker overnight. The separation and elution of IgG was done according to the manufacturer’s instructions. Eluted IgG was buffer exchanged into DMEM (VWR, 45000-306) using Amicon Ultra-4 Centrifugal Filter Unit, 30 kDa MWCO (Millipore Sigma, UFC803024) and concentrated to approximately 100 μL. After measuring the concentration, the purified IgG samples were stored at 4 °C.

#### Fusion peptide ELISA

Fusion peptide (FP) ELISA was performed as previously described^67^. Briefly, 384-well plates were coated with streptavidin and blocked with B3T buffer (150 mM NaCl, 50 mM Tris-HCl, 1 mM EDTA, 2% bovine serum albumin, 3.3% fetal bovine serum, 0.07% Tween 20, 0.02% thimerosal). After blocking, plates were incubated with FP-PEG12-biotin^39^. Serum sample was assessed in a 7-point, 4-fold serial dilution, followed by incubation with a 1:10,000 dilution of goat anti–guinea pig IgG conjugated to horseradish peroxidase (HRP) (SeraCare, Cat# 5220-0366). Plates were developed using 3,3′,5,5′-tetramethylbenzidine (TMB) and absorbance was measured at 450 nm. Nonlinear regression curves were fitted using a four-parameter asymmetric sigmoidal model (GraphPad Prism 10.4.2). EC1.5 (or ED1.5) values, defined as the concentrations (or dilution factors) yielding an OD450 of 1.5, were determined from the fitted curves.

#### HIV-1 Env trimer ELISA

HIV-1 Env trimer ELISA was conducted as described previously^67^. Briefly, 384-well plates were coated with Lectin and blocked with B3T buffer. After blocking, plates were incubated with BG505.DS.SOSIP trimer^39^. Serum was assessed in a 7-point, 4-fold serial dilution, followed by 10,000-fold dilution of goat anti-guinea pig IgG conjugated with horseradish peroxidase (HRP) (Sera care, Milford, MA, Cat#5220-0366). Plates were developed using 3,3’,5,5’-Tetramethylbenzidine (TMB) and read at 450 nM.

#### Neutralization

HIV-1 Env-pseudotyped virus stocks were generated by cotransfecting 293T cells with a pSG3ΔEnv backbone and an Env expression plasmid and neutralization assays were performed as previously described^67^. Briefly, serum samples were tested in 7-point, 4-fold serial dilutions starting at a 1:10 dilution, while control monoclonal antibodies (mAbs) were tested in 7-point, 5-fold dilutions starting at 50 µg/mL, using 384-well plates. For each well, 7.5 µL of diluted serum or mAb was mixed with 5 µL of HIV-1 Env pseudovirus and incubated for over 30 minutes at 37 °C. Next, 5 µL of TZM-bl cell suspension (2,500 cells per well) containing 0.07 mg/mL DEAE-Dextran hydrochloride was added to each well and incubated for 48 hours at 37 °C. On day 3, cells were lysed and incubated with Steadylite Plus (PerkinElmer, Waltham, MA) for 20 minutes at room temperature. Luminescence was measured using a luminometer to assess luciferase activity. A nonlinear regression dose-response curve was fitted using an asymmetric sigmoidal curve with 5-parameters. The 50% inhibitory concentrations (IC50 or ID50) were determined.

Neutralization with peptide competition was performed as described above, with the following modifications. A total of 3.75 µL of purified serum IgG was mixed with 3.75 µL of synthetic peptide, and the mixture was incubated at 37 °C for 30 minutes. Subsequently, 5 µL of HIV-1 Env pseudovirus was added. The final peptide concentration in the 12.5 µL reaction mixture was 50 µM. FP9v1 and its scrambled control peptide were synthesized (Genscript, Piscataway, NJ). All subsequent steps followed the standard neutralization assay protocol as described above.

The wildtype heterologous viruses were selected from a multi-clade panel of 208 HIV-1 strains using the following criteria: 1) the sequence of eight N-terminal amino acids (AVGIGAVF) of the fusion peptide matched the FP8.1 sequence; 2) the viral sequences showed a complete glycosylation profile surrounding the fusion peptide epitope. Specifically, N-linked glycosylation sites are observed at the amino acid positions of 88, 241, 448, 611, 625, and 637; 3) all strains are resistant to V3 and CD4i antibodies (17b, 48d, F105, 3074, 447-52D). Together, this is a multi-clade panel of strains with FP8.1 sequence, and it has excluded strains that are generally sensitive to V3 and CD4i antibodies or potentially super-sensitive to fusion peptide antibodies due to the lack of glycosylation.

#### Statistical Methods

Statistic analysis in this study was conducted in GraphPad Prism (Version 9.2.0).

