## Supplemental Figures for "Novel HIV-1 fusion peptide immunogens using glycan-engineered alphavirus-like particles"

#### Figure S1

| | CHIKV | VEEV | EEEV | WEEV |
|---|---|---|---|---|
| CHIKV | | 43% | 45% | 40% |
| VEEV | | | 54% | 46% |
| EEEV | | | | 48% |
| WEEV | | | | |

**Figure S1. Amino acid sequence identities between the E1 and E2 subunits of CHIKV, VEEV, EEEV and WEEV. Related to Figure 2.**

#### Figure S2

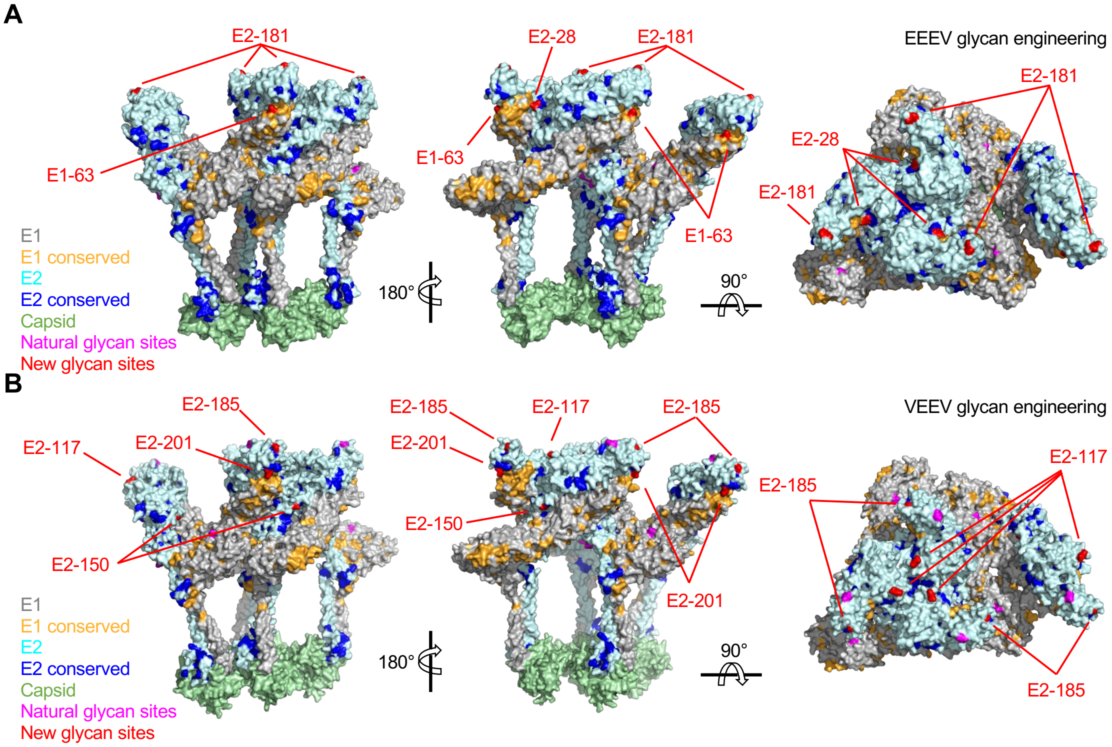


**Figure S2. Structure of the Capsid-E1-E2 of EEEV (PDB 6XO4, panel A) and VEEV (PDB 3J0C, panel B). Related to Figure 2.** Conserved residues across CHIKV, EEEV, and VEEV are highlighted. Preexisting and newly added glycan sites are highlighted.

### Figure S3

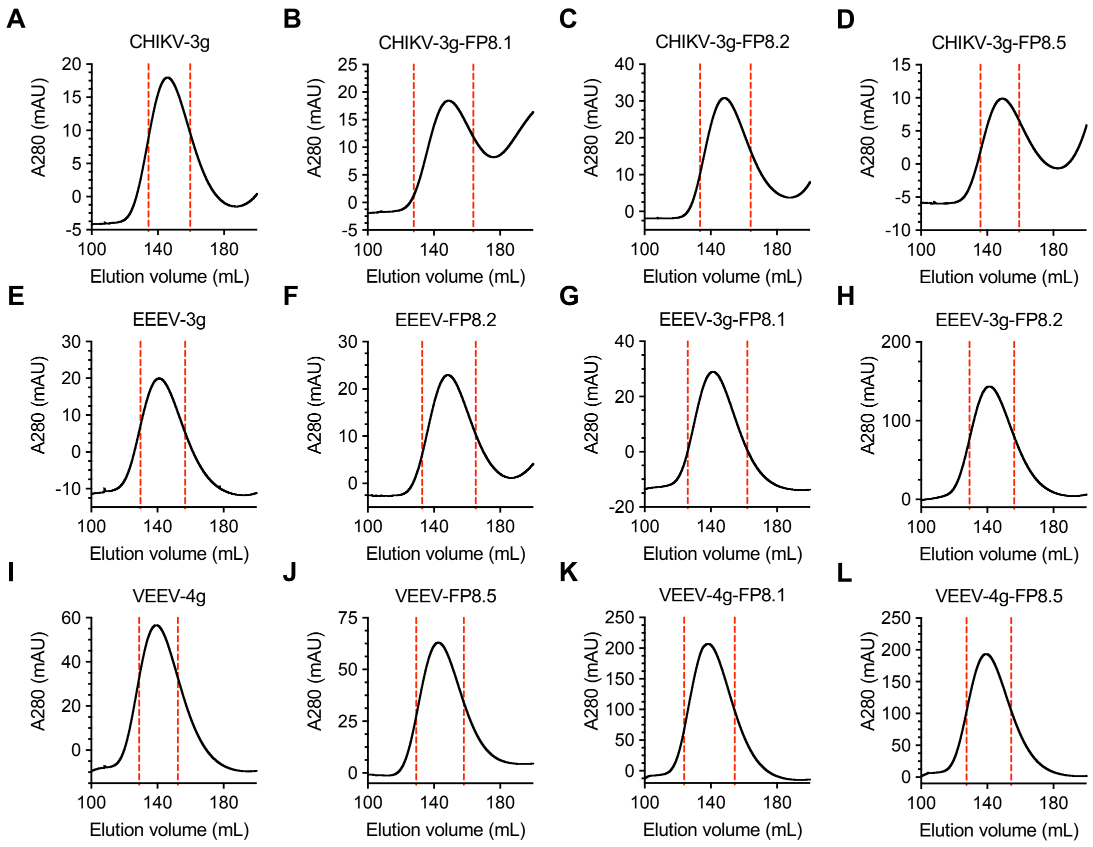


**Figure S3. VLP purification by size exclusion column. Related to Figure 2.**

### Figure S4

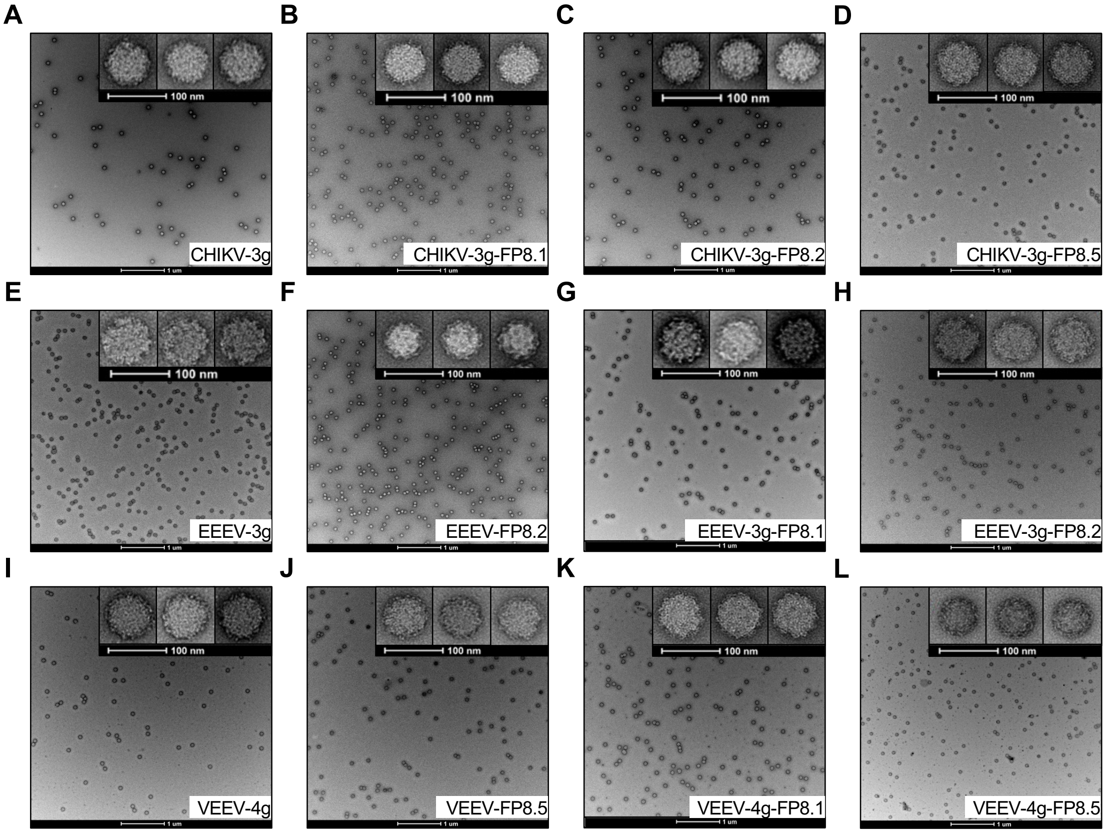


**Figure S4. VLP analysis by negative-stain electron microscopy.**

### Figure S5

| VLP | Glycan name | High-mannose | | Hybrid | | Complex | | Unoccupied | | Total Occupancy |
|---|---|---|---|---|---|---|---|---|---|---|
| | | Count | % | Count | % | Count | % | Count | % | % |
| CHIKV-3g-FP8.1 | E2-158 | 1383 | 35.0% | 858 | 21.7% | 1668 | 42.2% | 45 | 1.1% | 98.9% |
| | E1-69 | 0 | 0% | 0 | 0% | 886 | 87.9% | 122 | 12.1% | 87.9% |
| | E1-99 | 11 | 1.0% | 0 | 0% | 884 | 76.4% | 263 | 22.7% | 77.3% |
| EEEV-3g-FP8.1 | E2-28 | 1283 | 98.4% | 0 | 0% | 0 | 0% | 20 | 1.6% | 98.4% |
| | E2-181 | 70 | 1.0% | 45 | 0.6% | 7158 | 97.9% | 39 | 0.5% | 99.5% |
| | E1-63 | 350 | 18.4% | 247 | 13.0% | 1160 | 61.0% | 146 | 7.7% | 92.3% |
| EEEV-3g-FP8.2 | E2-28 | 2163 | 97.8% | 0 | 0% | 0 | 0% | 48 | 2.2% | 97.8% |
| | E2-181 | 130 | 1.5% | 2 | 0.0% | 8364 | 97.8% | 55 | 0.6% | 99.4% |
| | E1-63 | 477 | 16.9% | 303 | 10.7% | 1729 | 61.3% | 311 | 11.0% | 89.0% |
| VEEV-4g-FP8.1 | E2-117 | 36 | 0.7% | 92 | 1.8% | 4994 | 95.6% | 102 | 2.0% | 98.0% |
| | E2-150 | 3090 | 34.5% | 2803 | 31.3% | 2743 | 30.7% | 312 | 3.5% | 96.5% |
| | E2-185 | 6 | 8.8% | 4 | 5.8% | 55 | 80.3% | 3 | 5.1% | 94.9% |
| | E2-201 | 40 | 2.7% | 56 | 3.7% | 1297 | 86.1% | 113 | 7.5% | 92.5% |
| VEEV-4g-FP8.5 | E2-117 | 140 | 1.1% | 357 | 2.7% | 12031 | 92.6% | 460 | 3.5% | 96.5% |
| | E2-150 | 3315 | 32.1% | 3114 | 30.2% | 3432 | 33.3% | 452 | 4.4% | 95.6% |
| | E2-185 | 4 | 10.6% | 0 | 0% | 32 | 85.1% | 2 | 4.2% | 95.8% |
| | E2-201 | 25 | 1.8% | 43 | 3.2% | 1137 | 83.9% | 150 | 11.1% | 88.9% |

**Figure S5. Glycan analysis of purified VLPs. Related to Figure 2.**

### Figure S6

| Animal | BG505 | BI369.9A (clade A) | | | 286.36 (clade C) | | | CH117.4 (Clade BC) | | | BL01.DG (clade B) | | | SIVmac251.30 | | |
|---|---|---|---|---|---|---|---|---|---|---|---|---|---|---|---|---|
| code | No peptide | FP | Scr | %reduction | FP | Scr | %reduction | FP | Scr | %reduction | FP | Scr | %reduction | FP | Scr | %reduction |
| G5S1 | 0.0438 | >3.0 | 0.9 | >71% | >3.0 | >3.0 | No Neut | >3.0 | 1.0 | >65% | >3.0 | >3.0 | No Neut | >3.0 | >3.0 | No Neut |
| G5S2 | 0.0106 | >3.0 | 1.3 | >57% | >3.0 | >3.0 | No Neut | >3.0 | 0.6 | >81% | >3.0 | 2.1 | >31% | >3.0 | >3.0 | No Neut |
| G5S3 | 0.0908 | 3.8 | 0.9 | 77% | 4.2 | 3.8 | 8% | 3.1 | 0.9 | 71% | 3.4 | 2.4 | 28% | >5.8 | >5.8 | No Neut |
| G5S4 | 0.3772 | 3.0 | 0.9 | 69% | >3.9 | >3.9 | No Neut | >3.9 | 0.5 | >87% | >3.9 | 3.7 | >6% | >3.9 | >3.9 | No Neut |
| G5S5 | 0.4447 | 4.6 | 1.3 | 72% | >5.8 | >5.8 | No Neut | >5.8 | 0.8 | >87% | >5.8 | 2.5 | >58% | >5.8 | >5.8 | No Neut |
| G5S6 | 0.0078 | 2.7 | 0.9 | 66% | >4.7 | >4.7 | No Neut | >4.7 | 0.4 | >91% | >4.7 | 2.0 | >58% | >4.7 | >4.7 | No Neut |
| G5S7 | 0.1034 | >4.3 | 2.1 | >51% | >4.3 | >4.3 | No Neut | 3.6 | 2.3 | 38% | 2.4 | 1.3 | 44% | >4.3 | >4.3 | No Neut |
| G5S8 | 0.1024 | 5.1 | 1.5 | 71% | 5.4 | >5.4 | No Neut | >5.4 | 0.8 | >85% | 5.1 | 2.3 | 56% | >5.4 | >5.4 | No Neut |
| G5S25 | 0.3496 | >4.6 | 3.6 | >22% | >4.6 | >4.6 | No Neut | >4.6 | 3.0 | >35% | >4.6 | >4.6 | No Neut | >4.6 | >4.6 | No Neut |
| G5S26 | 0.1190 | >7.5 | 3.3 | >57% | >7.5 | >7.5 | No Neut | >7.5 | 3.3 | >56% | >7.5 | 6.6 | >12% | >7.5 | >7.5 | No Neut |
| G5S27 | 0.0018 | >5.1 | 3.3 | >34% | 1.6 | 1.4 | 11% | 1.4 | 0.7 | 49% | >5.1 | >5.1 | No Neut | >5.1 | >5.1 | No Neut |
| G5S28 | 0.0039 | >5.2 | 2.9 | >44% | >5.2 | >5.2 | No Neut | 4.1 | 2.8 | 33% | 4.9 | 3.8 | 23% | >5.2 | >5.2 | No Neut |
| G5S29 | 0.1995 | 5.3 | 0.4 | 93% | 2.5 | 1.8 | 27% | 2.9 | 0.3 | 90% | 2.9 | 0.6 | 78% | >5.9 | >5.9 | No Neut |
| G5S30 | 0.0822 | >7.1 | 1.0 | >85% | >7.1 | 5.3 | >26% | 6.7 | 0.8 | 89% | 3.0 | 1.9 | 38% | >7.1 | >7.1 | No Neut |
| G6S9 | 0.0007 | 3.2 | 1.4 | 57% | 2.2 | 2.2 | 1% | 2.9 | 1.1 | 61% | 2.5 | 1.7 | 31% | >4.8 | >4.8 | No Neut |
| G6S10 | 0.0873 | >7.6 | 1.8 | >77% | >7.6 | >7.6 | No Neut | 4.1 | 1.0 | 76% | 3.1 | 2.0 | 38% | >7.6 | >7.6 | No Neut |
| G6S11 | 0.0557 | >4.0 | 3.2 | >19% | >4.0 | >4.0 | No Neut | >4.0 | 1.9 | >52% | >4.0 | 2.8 | >31% | >4.0 | >4.0 | No Neut |
| G6S12 | 0.0022 | >4.1 | >4.1 | No Neut | >4.1 | >4.1 | No Neut | >4.1 | >4.1 | No Neut | >4.1 | 4.0 | >2% | >4.1 | >4.1 | No Neut |
| G6S13 | >1.676 | 2.7 | 0.9 | 66% | >5.0 | >5.0 | No Neut | >5.0 | 0.6 | >88% | >5.0 | 2.4 | >52% | >5.0 | >5.0 | No Neut |
| G6S14 | 0.0066 | >4.5 | 3.0 | >33% | >4.5 | >4.5 | No Neut | >4.5 | 3.2 | >29% | >4.5 | >4.5 | No Neut | >4.5 | >4.5 | No Neut |
| G6S15 | 0.1329 | >3.7 | >3.7 | No Neut | >3.7 | >3.7 | No Neut | >3.7 | >3.7 | No Neut | >3.7 | >3.7 | No Neut | >3.7 | >3.7 | No Neut |
| G6S16 | 0.0005 | >6.6 | >6.6 | No Neut | >6.6 | >6.6 | No Neut | >6.6 | >6.6 | No Neut | >6.6 | >6.6 | No Neut | >6.6 | >6.6 | No Neut |
| G7S17 | 0.2219 | 3.9 | 0.4 | 90% | >7.6 | 6.1 | >19% | 5.3 | 0.7 | 87% | 5.9 | 1.8 | 69% | >7.6 | >7.6 | No Neut |
| G7S18 | 0.2160 | 2.7 | 0.4 | 85% | >3.0 | >3.0 | No Neut | >3.0 | 0.7 | >75% | >3.0 | 1.8 | >40% | >3.0 | >3.0 | No Neut |
| G7S19 | 0.0853 | >3 | >3 | No Neut | >3.0 | >3.0 | No Neut | >3.0 | 2.2 | >26% | >3.0 | >3.0 | No Neut | >3.0 | >3.0 | No Neut |
| G7S20 | 0.1562 | >4.2 | 2.6 | >37% | 1.6 | 1.4 | 14% | 1.0 | 0.4 | 57% | >4.2 | >4.2 | No Neut | >4.2 | >4.2 | No Neut |
| G7S21 | 0.0602 | >4.3 | 1.0 | >77% | >4.3 | >4.3 | No Neut | >4.3 | 1.1 | >74% | >4.3 | 3.6 | >17% | >4.3 | >4.3 | No Neut |
| G7S22 | 0.1355 | >3.4 | 2.2 | >36% | >3.4 | >3.4 | No Neut | >3.4 | 1.5 | >57% | >3.4 | 3.0 | >12% | >3.4 | >3.4 | No Neut |
| G7S23 | 0.0556 | 1.6 | 0.3 | 81% | 3.8 | 2.5 | 35% | 3.7 | 0.2 | 95% | 1.4 | 1.0 | 29% | >6.6 | >6.6 | No Neut |
| G7S24 | 0.0245 | 2.5 | 0.7 | 72% | >4.5 | >4.5 | No Neut | 1.5 | 0.4 | 73% | 1.2 | 0.7 | 41% | >4.5 | >4.5 | No Neut |

**Figure S6. Neutralization IC50 titers of purified guinea pig serum IgG with or without the presence of free FP peptide. Related to Figure 6.** IC50 titers are shown in mg/mL. Animal code GxSy indicates Group x subject y. When 50% neutralization is not achieved at highest tested concentration, the IC50 titer is shown as ">highest tested concentration". FP and scr indicate neutralization in the presence of fusion peptide or scramble peptide, respectively. %reduction is calculated as 1-(IC50[src]/IC50[FP]). %reduction greater than 30% is highlighted in orange, indicating the presence of FP-directed neutralization. %reduction that is potentially greater than 30% is highlighted in green, indicating the potential presence of FP-directed neutralization.
